# Inhibition of a negative feedback for persistent epithelial cell–cell junction contraction by p21-activated kinase 3

**DOI:** 10.1101/743237

**Authors:** Hiroyuki Uechi, Erina Kuranaga

## Abstract

Actin-mediated mechanical forces are central drivers of cellular dynamics. They generate protrusive and contractile dynamics, the latter of which are induced in concert with myosin II bundled at the site of contraction. These dynamics emerge concomitantly in tissues and even each cell; thus, the tight regulation of such bidirectional forces is important for proper cellular deformation. Here, we show that contractile dynamics can eventually disturb cell–cell junction contraction in the absence of p21-activated kinase 3 (Pak3). Upon Pak3 depletion, contractility induces the formation of abnormal actin protrusions at the shortening junctions, which reduces E-cadherin levels at adherens junctions. Such E-cadherin dilution dissociates myosin II from the contracting junctions, leading to a reduction in junctional tensile forces. Overexpressing E-cadherin restores the association of myosin II at the junctions and junction contraction. Our results suggest that contractility both induces and perturbs junction contraction and that the attenuation of such perturbations by Pak3 facilitates persistent junction shortening.

---

The cell collectives composing animal bodies sculpt tissue architectures through various cellular behaviors such as cell division, deformation, rearrangement, and migration. The cytoskeletal protein actin is the central protein that drives these cellular behaviors by generating mechanical forces^1–7^. While actin generates protrusive forces by forming branched and bundled structures, it also supplies contractile forces by forming bundled structures and loosely organized networks in concert with non-muscle myosin II. Such bidirectional force generation by actin can coexist in each cell and even at the same position within cells. During single-cell migration, protrusive actin dynamics extend cells at the leading edge, while actomyosin (the actin and myosin II complex) contraction causes retraction at the rear of the cell along the migrating direction^8, 9^. A recent study of *Drosophila* eye development demonstrated that pulsatile extension by protrusive branched actin networks and counterbalancing actomyosin contractility-mediated shortening of each cell–cell contact control cellular shape^10^. Thus, the tight regulation of actin dynamics is important for proper force induction and the resultant cellular dynamics.

Epithelial cell intercalation is one of the multicellular dynamics driven by the contractile forces of actomyosin and contributes to directional tissue extension and movement^11–13^. This process consists of the directional exchange of cellular positions within cell collectives, which is driven by cell–cell junction remodeling (i.e., shortening of cell–cell junctions and subsequent growth of new ones in new directions)^14^. During shortening, actin and myosin II are highly enriched at the adherens junctions (AJs) of cell–cell junctions to form contractile actomyosin bundles that then shorten the junctions^1, 11, 15–19^. The mechanisms inducing contractile forces via actomyosin are well studied; however, it is still unknown whether actomyosin-mediated contractions are negatively affected during cell–cell junction shortening, and if so, how the shortening is sustained.

The rotation of *Drosophila* male genitalia is an example of epithelial cell intercalation^20^. During metamorphosis, male genitalia located at the posterior end of the body undergo 360° dextral rotation around the anterior–posterior axis (Fig. 1a)^21, 22^. This rotation is observed from 24 to 36 h after puparium formation (APF), and is composed of two movements of epithelial cells: the initial 180° dextral movement of the genitalia along with the surrounding epithelia, which is called the posterior compartment of the A8 segment (A8p), at the anterior side of the genitalia, and the subsequent 180° dextral movement of the anterior compartment of the A8 segment (A8a), the latter of which starts at 26 h APF^20–22^. During rotation, the A8a cells frequently induce left–right polarized junction remodeling in relation to the anterior– posterior axis in the confined space between the A8p epithelia and the A7 segment, which results in unidirectional epithelial cell movement^20, 23^. Consistent with other tissues, the accumulation of actomyosin at the AJs of cell–cell junctions induces junction contraction in this model. Down-regulation of contractile activity, such as via RNA interference (RNAi) of myosin II regulatory light chain (MRLC), compromises the remodeling and hence results in insufficient A8a cell movement, leading to incomplete rotation of the genitalia^20^.

**Figure 1.**
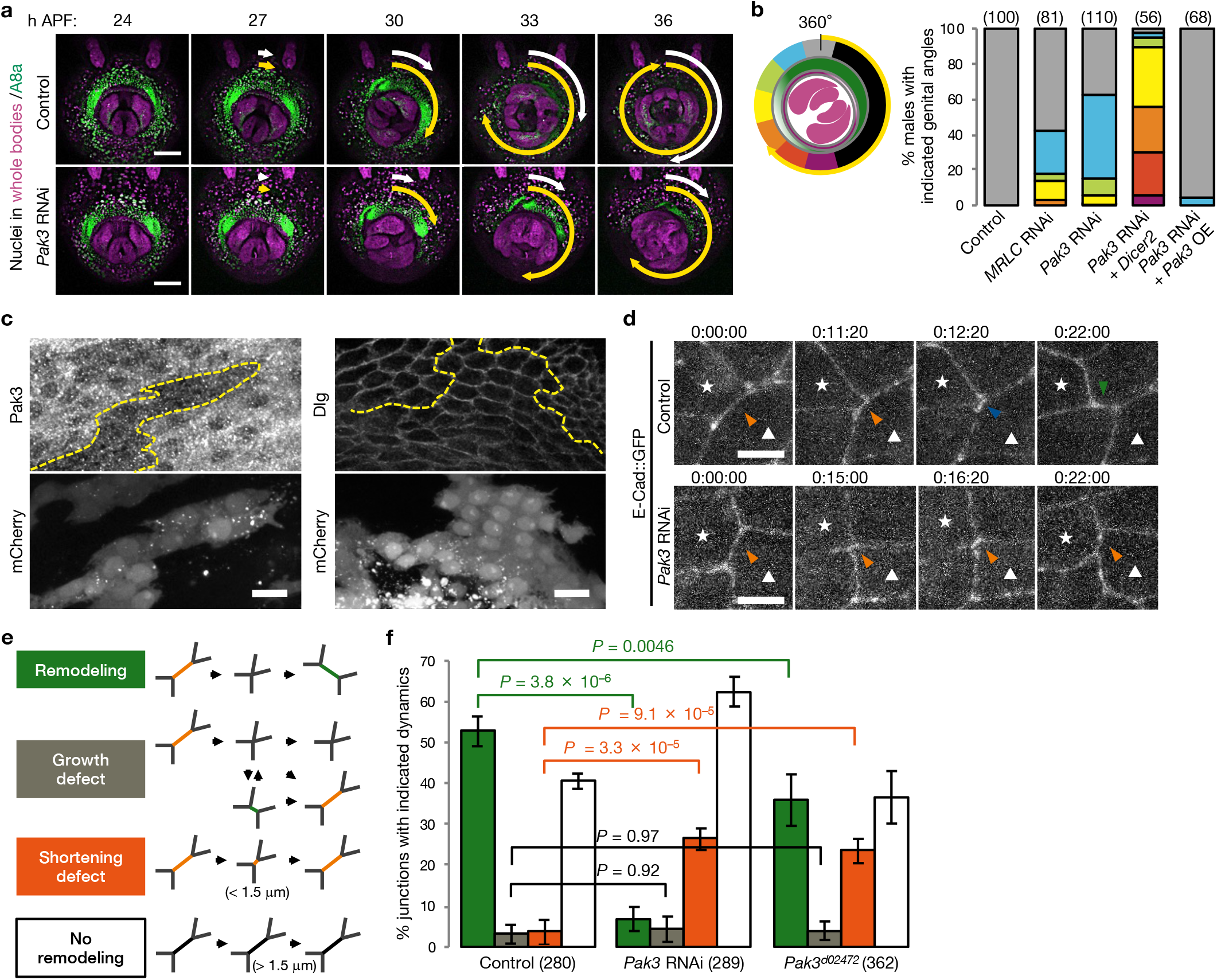
Pak3 is required for cell–cell junction shortening during epithelial junction remodeling. (**a**) Representative time-lapse images of male genitalia observed from the posterior end during rotation. The ventral side is located at the top. Nuclei in the A8a cells and the whole body are depicted in green and purple, respectively. Yellow and white arrows indicate the movements of genitalia and the A8a epithelia, respectively. Scale bar, 100 μm. (**b**) Left, schema for the categorization of genitalia angles. Yellow arrow indicates genitalia movement. Gray indicates the 360° rotation of genitalia (normal orientation). Right, percentages of male adult flies with the indicated genitalia angles are shown. Parentheses indicate the number of males examined. (**c**) A8a epithelia at 28 h APF immunostained for Pak3 and Dlg. Pak3 RNAi cells are indicated with mCherry signals. Scale bar, 10 μm. Broken lines indicate the edges of Pak3 RNAi clones. (**d**) Representative time-lapse images of E-Cad::GFP at remodeling junctions. Stars and triangles indicate cells forming the shortening junctions. Orange, blue, and green arrowheads indicate shortening junctions, four-way vertices, and growing junctions, respectively. Scale bar, 5 μm. (**e**) Schema representing the categorization of junction dynamics. (**f**) Mean ± S.D. of the percentages of junctions with the categorized dynamics. Bar colors correspond to (e). Parentheses indicate the number of examined junctions from 3 pupae per genotype. *P-*values by Tukey’s test. Genotypes: (**a**) *+/Y;His2Av::mRFP/+;AbdB-Gal4, UAS-H2B::ECFP/+* (Control) and *+/Y;His2Av::mRFP/+;AbdB-Gal4, UAS-H2B::ECFP/UAS-Pak3 RNAi*; (**b**) *+/Y;;AbdB-Gal4/+* (Control), *+/Y;;AbdB-Gal4/UAS-MRLC RNAi*, *+/Y;;AbdB-Gal4/UAS-Pak3 RNAi*, *+/Y;UAS-Dicer2/+;AbdB-Gal4/UAS-Pak3 RNAi*, and *+/Y;;AbdB-Gal4, UAS-Pak3::GFP/UAS-Pak3 RNAi*; (**c**) *hs-flp/Y;E-Cad::GFP;Act>CD2>Gal4, UAS-mCD8::mCherry/UAS-Pak3 RNAi*; (**d** and **f**) *+/Y;E-Cad::GFP;AbdB-Gal4, UAS-H2B::ECFP/+* (Control), *+/Y;E-Cad::GFP;AbdB-Gal4, UAS-H2B::ECFP/UAS-Pak3 RNAi*, and *+/Y;E-Cad::GFP;Pak3^d02472^*.

Here, we examined actin and myosin II dynamics in the A8a epithelia during the rotation of *Drosophila* genitalia and revealed that, upon junction contraction, actin dynamics at cell–cell junctions are compromised in p21-activated kinase 3 (Pak3) mutant flies. These aberrant actin dynamics disturb the distribution of E-cadherin and myosin II at the junctions, and thus eventually disrupt the junction contraction. These findings suggest that Pak3 blocks the negative feedback of contractility and ensures persistent junction contraction and rearrangement of epithelial cells.

## Results

### Pak3 is required for A8a cell movement during the rotation of *Drosophila* genitalia

To understand the molecular basis underlying epithelial cell–cell junction remodeling, we searched for genes involved in the movement of A8a cells. In the control flies, time-lapse images showed that male genitalia underwent the 360° full rotation during metamorphosis, and all male flies had the normal orientation of genitalia (Fig. 1a,b). We found that RNAi targeting Pak3 in the A8a epithelia under the control of the *AbdB-Gal4* driver caused insufficient movement of the A8a epithelia during rotation, while the genitalia still underwent the 180° movement with respect to the A8a epithelia (Fig. 1a)^24, 25^. In addition, depleting Pak3 induced the misorientation of genitalia in approximately 60% of adult male flies, similar to that reported for MRLC depletion (Fig. 1b)^20^. Co-expression of Dicer2, which augments RNAi efficiency^26^, with Pak3 RNAi increased the frequency of misorientation to > 90%, which was associated with a decline in rotation angle, while the overexpression (OE) of Pak3 tagged with green fluorescent protein (Pak3::GFP) restored the normal orientation, excluding the possibility of an off-target effect of this Pak3 RNAi (Fig. 1b). These results indicate that Pak3 is involved in epithelial tissue dynamics.

Immunostaining of fixed epithelia revealed that Pak3 was expressed in the A8a cells, as indicated by its decreased signals in Pak3 RNAi cells that were clonally introduced into wild-type tissues (Fig. 1c). Signals for the lateral membrane-associating protein Discs large (Dlg)^27^ at cell–cell boundaries were indistinguishable between wild-type cells and Pak3 RNAi clones, suggesting that Pak3 RNAi does not compromise the integrity of epithelial cells (Fig. 1c).

### Pak3 is required for cell–cell junction shortening

To examine the role of Pak3 in cellular dynamics, we monitored the dynamics of cell–cell junctions labeled with GFP-tagged E-cadherin (E-Cad::GFP), an adhesive AJ component, during A8a cell movement^28, 29^. In the control epithelia, we observed that cell–cell junctions undergo shortening, form four-way vertices, and subsequently grow in other directions, consistent with a previous report (Fig. 1d; Supplementary Movie 1)^20^. In the Pak3-depleted A8a cells, junctions frequently failed to complete shortening; that is, they did not shorten sufficiently to form four-way vertices and instead went back to the original direction (Fig. 1d; Supplementary Movie 2). To quantify these defective junction dynamics, we categorized the junctions in A8a cells according to their dynamics as follows: junctions that completed shortening and grew in other directions (remodeling), junctions that completed shortening, but failed to grow in other directions after forming four-way vertices (growth defect), junctions that shortened sufficiently to reach 1.5 μm in length, but failed to form four-way vertices (shortening defect), and junctions that did not shorten to less than 1.5 μm in length (no remodeling) (Fig. 1e). Then, the populations of junctions in each category were examined. In the control cells, approximately 55% of junctions underwent remodeling, and few junctions showed defects during a 4-h period following the initiation of A8a cell movement (from 26 to 30 h APF, Fig. 1f). By contrast, in Pak3 RNAi cells, the frequency of junction remodeling was decreased to < 10%, and instead, the number of shortening junctions that failed to form four-way vertices (categorized as a shortening defect) was significantly increased (Fig. 1f). Similar propensities were observed in the A8a cells of a Pak3 hypomorphic mutant (*Pak3^d02472^*)^30^, although the decline in remodeling frequency was not substantial compared to that in Pak3 RNAi cells (Fig. 1f). Meanwhile, the frequency of the growth defect was not altered in these Pak3 mutants (Fig. 1f). These results suggest that Pak3 is required for junction shortening during cell intercalation.

### Pak3 suppresses aberrant actin dynamics in response to junction contraction

Junction shortening is driven by the contractile forces of actomyosin accumulating at the AJs of junctions, and Pak protein families are known to regulate actin dynamics^5, 31–33^. These findings and our observations led us to speculate that Pak3 has roles in actin dynamics during junction shortening in the A8a epithelia. To test this possibility, we performed time-lapse imaging of GFP-tagged Lifeact (Lifeact::GFP), which labels F-actin^34^. In the control A8a cells, actin was distributed predominantly at cell–cell junctions (Fig. 2a; Supplementary Movie 3). Magnified images showed that, whereas the majority of Lifeact::GFP signals were localized along junctions, actin frequently generated small protrusions arising from the junctions (Fig. 2b). The maximum size of these actin-positive protrusions was 1.1 ± 0.34 μm in height (perpendicular to the junctions) and 2.1 ± 1.1 μm in width (parallel to the junctions) (Fig. 2c). These observations suggest that actin not only merely forms into bundled structures along cell–cell junctions but also generates small protrusive structures in the A8a cells undergoing junction remodeling. In Pak3 RNAi cells, actin also localized to cell–cell junctions, but generated significantly larger protrusive structures, compared to those in the control cells (Fig. 2a–c; Supplementary Movie 4). Similar actin-positive structures were also observed in *Pak3^d02472^* flies with another actin-labeling probe, GFP-tagged actin-binding domain of utrophin (UtrABD::GFP) (Fig. 2d)^35, 36^. To semi-quantify these actin dynamics in Pak3 RNAi cells, we defined a “large” actin protrusion as a Lifeact-positive cluster at junctions that was greater than 1.5 μm in height and greater than 3 μm in width in a planar section; both lengths approximately exceeded the mean + 1 standard deviation (S.D.) of each length of the actin protrusions in the control cells, respectively (Fig. 2c,e). We then examined the frequency of the appearance of such large protrusions at each junction (Fig. 2f). While greater than 95% of cell–cell junctions in the control cells did not generate large protrusions, these structures emerged at almost all junctions at least once per hour in Pak3 RNAi cells (Fig. 2f). Again, overexpressing Pak3 partially alleviated the formation of large protrusions: although these structures were still observed in approximately 50% of junctions, they were absent from greater than 40% of junctions, and only a few junctions formed large protrusions more than three times per hour (Fig. 2f). These data suggest that Pak3 suppresses the formation of aberrant actin protrusions in the A8a cells.

**Figure 2.**
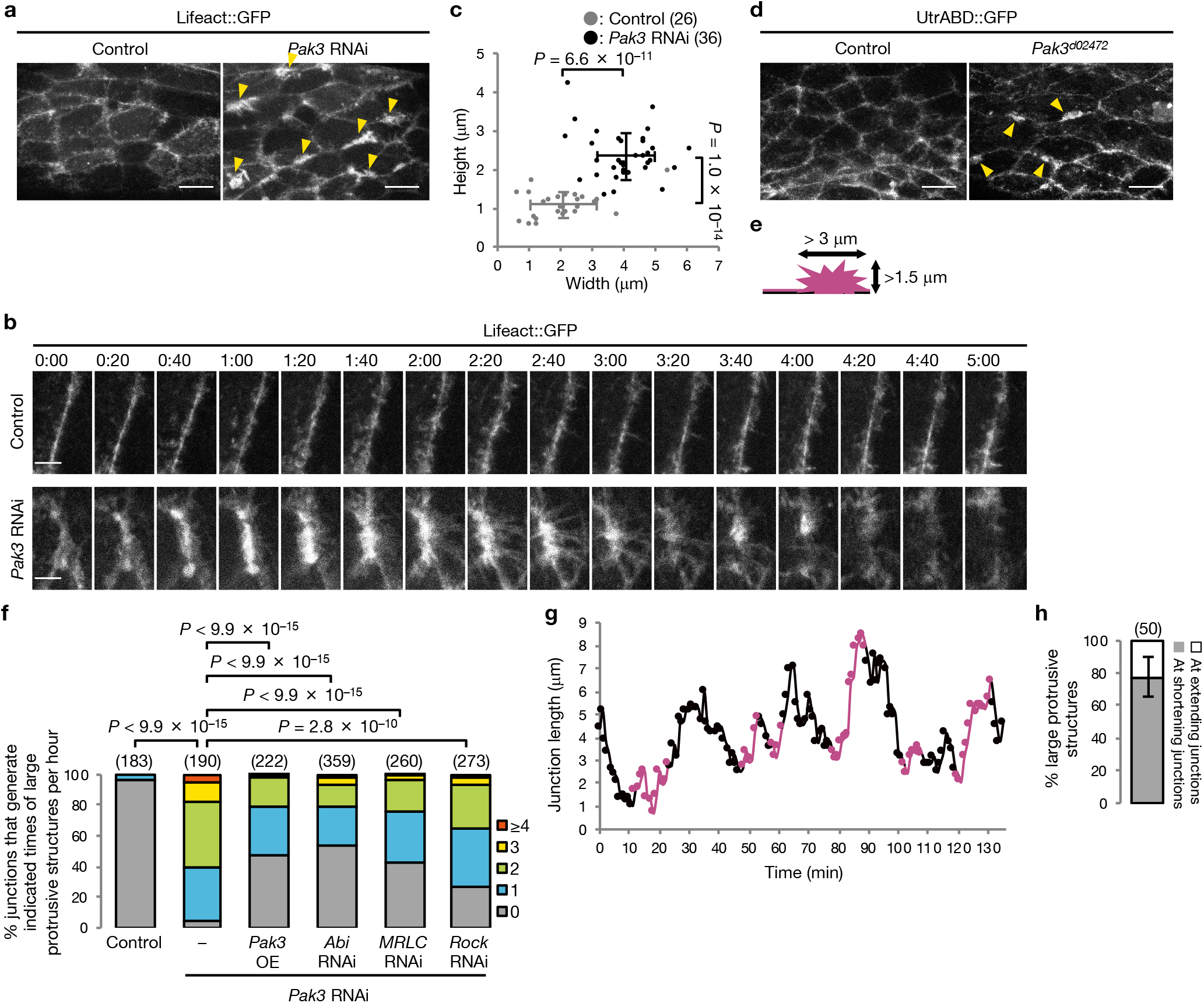
Pak3 depletion enhances the formation of large actin protrusions. (**a**) Images of actin labeled with Lifeact::GFP in the A8a cells. Arrowheads indicate some actin protrusions. Scale bar, 10 μm. (**b**) Magnified time-lapse images of Lifeact::GFP at cell–cell junctions. Scale bar, 3 μm. (**c**) Mean ± S.D. of the width and height of actin protrusion. Parentheses indicate the number of examined protrusions from 5 pupae per genotype. *P-*values by unpaired *t*-test. (**d**) Images of actin labeled with UtrABD::GFP in the A8a cells. Arrowheads indicate some actin protrusions. Scale bar, 10 μm. (**e**) Schema of large actin (magenta) protrusions at a boundary (black). (**f**) Percentages of junctions generating large actin protrusions for the indicated number of times per hour are shown. Parentheses indicate the number of examined junctions from 4–5 pupae per genotype. *P-*values by Mann-Whitney *U*-test. (**g**) Graph showing the length of a representative junction that repeats shortening and re-extension. The timings when large actin protrusions emerge are shown in magenta. (**h**) Mean ± S.D. of the timing of the onset of large actin protrusion formation. Parenthesis indicates the number of protrusions from 15 junctions of 3 pupae. Genotypes: (**a**–**c**) *+/Y;UAS-Lifeact::GFP/+;AbdB-Gal4/+* (Control) and *+/Y;UAS-Lifeact::GFP/+;AbdB-Gal4/UAS-Pak3 RNAi*; (**d**) *+/Y;sqh-UtrABD::GFP/+* (Control) and *+/Y;sqh-UtrABD::GFP/+;Pak3^d02472^*; (**f**) *+/Y;UAS-Lifeact::GFP/+;AbdB-Gal4/+* (Control), *+/Y;UAS-Lifeact::GFP/+;AbdB-Gal4/UAS-Pak3 RNAi*, *+/Y;UAS-Lifeact::GFP/+;AbdB-Gal4, UAS-Pak3::GFP/UAS-Pak3 RNAi*, *+/Y;UAS-Lifeact::GFP/UAS-Abi RNAi;AbdB-Gal4/UAS-Pak3 RNAi*, *+/Y;UAS-Lifeact::GFP/UAS-MRLC RNAi;AbdB-Gal4/UAS-Pak3 RNAi*, and *+/Y;UAS-Lifeact::GFP/UAS-Rock RNAi;AbdB-Gal4/UAS-Pak3 RNAi*; (**g** and **h**) *+/Y;UAS-Lifeact::GFP/+;AbdB-Gal4/UAS-Pak3 RNAi*.

Branched actin networks induce the formation of protrusive actin structures and are generated by the Arp2/3 complex, which is stimulated by the WAVE regulatory complex (WRC)^1, 3^. To determine whether this pathway is involved in the formation of large protrusions, we used RNAi targeting Abi, a component of the WRC, in Pak3-depleted cells^37, 38^. Depletion of Abi suppressed the generation of large actin protrusions, although it did not completely restore actin dynamics (Fig. 2f). This suggests that the aberrant actin protrusions are in part composed of branched actin-networks.

To further characterize these aberrant actin protrusions in Pak3 RNAi cells, we explored the correlation between their emergence and junction dynamics. In Pak3-depleted cells, junctions failed to undergo remodeling (Fig. 1f), but instead underwent repeated shrinkage and extension (Fig. 2g). We found that the generation of large protrusions was initiated frequently when the junctions were shortening rather than when they were extending (Fig. 2g). Approximately 80% of the large actin protrusions emerged at shortening junctions (Fig. 2h). These observations raise the possibility that junction contraction sensitizes cells to the formation of large protrusions. To evaluate this possibility, we depleted MRLC or Rock, an upstream activator of myosin II^1, 5^. Depletion of these factors suppressed the emergence of large protrusions in Pak3 RNAi cells (Fig. 2f). Collectively, these results suggest that Pak3 suppresses the formation of branched network-containing actin protrusions at cell–cell junctions upon contraction of the junctions.

### Pak3 retains myosin II cables and tension at cell–cell junctions

During the emergence of aberrant actin protrusions, the junctions failed to continue shortening and instead re-extended (Fig. 2g), implying that while large protrusions are induced upon junction contraction, they in turn perturb contraction. To understand how junction contraction is compromised, we examined the dynamics of myosin II. Time-lapse imaging of GFP-tagged MRLC (MRLC::GFP) showed that the majority of MRLC::GFP signals were observed as a single cable at cell–cell junctions (Fig. 3a). In the control cells, the MRLC::GFP signals still remained as a single cable at the initiation of junction shortening; however, when the junction lengths were decreased, the MRLC::GFP signals split into two distinct cables at the shortening junctions (Fig. 3a, arrowheads). In Pak3-depleted cells, we found that the MRLC::GFP cables had already split before the junctions had shortened sufficiently (Fig. 3a, arrowheads). Measurement of the lengths of junctions when MRLC::GFP cables split revealed that depleting Pak3 significantly increased their length from 1.3 ± 0.41 to 2.2 ± 0.72 μm (Fig. 3b). Since E-Cad::GFP signals did not segregate at shortening junctions (Fig. 1d), this observation suggests that myosin II is dissociated from shortening cell–cell junctions.

**Figure 3.**
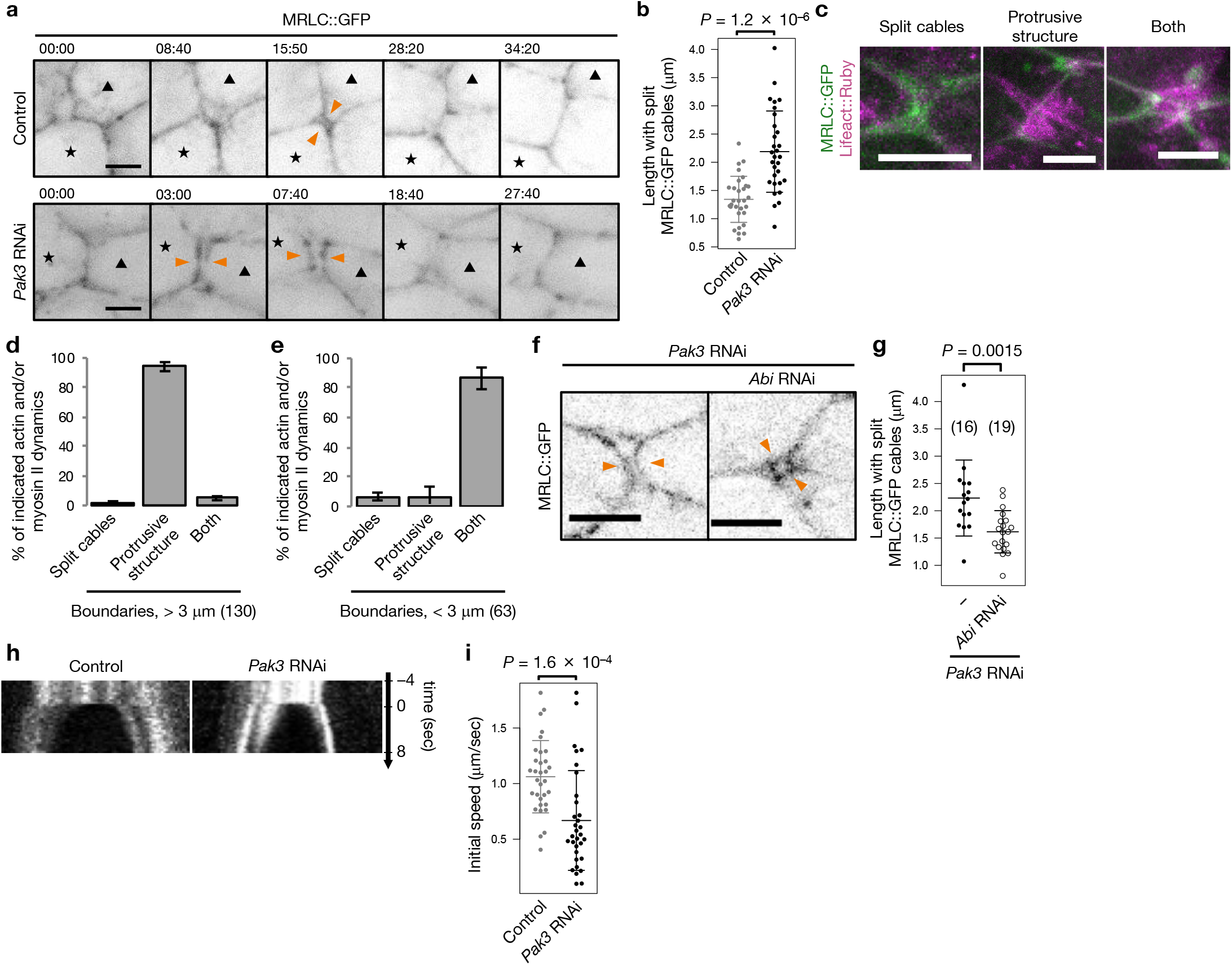
Pak3 depletion causes the dissociation of myosin II and reduces tension at junctions. (**a**) Time-lapse images of MRLC::GFP at shortening junctions. Stars and triangles indicate cells forming the shortening junctions. Orange arrowheads indicate splitting myosin II cables. Scale bar, 3 μm. (**b**) Mean ± S.D. of the lengths of junctions with split myosin II cables; 30 junctions from 3 (Control) and 4 (Pak3 RNAi) pupae were examined. *P*-value by unpaired *t*-test. (**c**) Representative images of cell–cell junctions with split myosin II cables, large actin protrusions, and both (from the left). Scale bar, 5 μm. (**d** and **e**) Mean ± S.D. of the percentages of aberrant actin and myosin II dynamics as categorized in (c) at cell–cell junctions larger than (d) and smaller than (e) 3 μm. Parentheses indicate the number of examined junctions from 3 pupae. (**f**) Representative images of split MRLC::GFP cables (orange arrowheads) at junctions. Scale bar, 5 μm. (**g**) Mean ± S.D. of the lengths of junctions with split myosin II cables. Parentheses indicate the number of examined junctions from 5 (Pak3 RNAi) and 6 (Pak3 and Abi RNAis) pupae. *P*-value by Dunnett’s test. (**h**) Kymographs of cell– cell junctions labeled with E-Cad::GFP and ablated with a 365-nm laser at *t* = 0. (**i**) Mean ± S.D. of the initial speed of vertex displacement after ablation; 32 junctions from 9 (Control) and 8 (Pak3 RNAi) pupae were examined. *P-*value by unpaired *t*-test. Genotypes: (**a** and **b**) *MRLC^AX3^/Y;MRLC-MRLC::GFP;AbdB-Gal4, UAS-H2B::ECFP/+* (Control) and *MRLC^AX3^/Y;MRLC-MRLC::GFP;AbdB-Gal4, UAS-H2B::ECFP/UAS-Pak3 RNAi*; (**c**–**e**) *MRLC^AX3^/Y;MRLC-MRLC::GFP/UAS-Lifeact::Ruby;AbdB-Gal4, UAS-H2B::ECFP/UAS-Pak3 RNAi*; (**f** and **g**) *MRLC^AX3^/Y;MRLC-MRLC::GFP/+;AbdB-Gal4, UAS-H2B::ECFP/UAS-Pak3 RNAi* and *MRLC^AX3^/Y;MRLC-MRLC::GFP/UAS-Abi RNAi;AbdB-Gal4, UAS-H2B::ECFP/UAS-Pak3 RNAi*; (**h** and **i**) *+/Y;E-Cad::GFP;AbdB-Gal4, UAS-H2B::ECFP/+* (Control) and *+/Y;E-Cad::GFP;AbdB-Gal4, UAS-H2B::ECFP/UAS-Pak3 RNAi*.

To examine the interplay between the large actin protrusions and myosin II cables in Pak3 RNAi cells, we next observed myosin II and actin dynamics simultaneously using MRLC::GFP and Lifeact tagged with red fluorescent protein (Lifeact::Ruby)^34^. We categorized the aberrant actin and myosin II dynamics into the following three groups and examined their proportions: splitting of myosin II cables, generation of large actin protrusions, and both (Fig. 3c–e). At junctions greater than 3 μm in length, which roughly exceed the mean + 1 S.D. of the length with split myosin II cables in Pak3-depleted cells (Fig. 3b), large protrusive structures were still generated in the absence of myosin II splitting, suggesting that the dissociation of myosin II from cell–cell junctions is not located upstream from the formation of aberrant actin protrusions (Fig. 3d). In contrast, at junctions less than 3 μm in length (or shortening junctions), most of the splitting cables were observed concomitantly with large actin protrusions (Fig. 3e). This observation raises the possibility that the emergence of large actin protrusions triggers the segregation of myosin II at shortening junctions. To test this possibility, we depleted Abi in Pak3 RNAi cells. This manipulation reduced the length at which the junctions showed split myosin II cables, suggesting that myosin II dissociation is associated with the generation of aberrant actin protrusions (Fig. 3f,g).

We hypothesized that the failure of junction contraction observed in Pak3-depleted cells was attributed to the dissociation of myosin II from shortening junctions and the consequent reduction in tension at the junctions. To evaluate tension at junctions, we cut the junctions by ablation with a 365-nm laser and measured the displacement of their vertices, which reflects junctional tension (Fig. 3h,i)^39, 40^. This showed that the initial speed of displacement upon laser ablation was decreased in Pak3 RNAi cells (Fig. 3i). Collectively, these results suggest that Pak3 ensures junction contraction by retaining myosin II attachment and tension at cell–cell junctions.

### Dilution of E-cadherin at cell–cell junctions mediates myosin II dissociation

To clarify how the large actin protrusions and consequent segregation of myosin II at shortening junctions are connected, we again explored the distribution of E-cadherin, since it is a central component of the cadherin–catenin core complex that acts as a scaffold for actomyosin at AJs^29, 41^. Magnified time-lapse images of E-Cad::GFP in the control cells revealed that while the majority of E-Cad::GFP signals were localized along cell–cell junctions, they frequently formed small protrusions arising from the junctions (Fig. 4a, arrowheads; Supplementary Movie 5). These protrusions were positive for E-Cad::GFP and Lifeact::Ruby, suggesting that E-cadherin is associated with actin protrusions (Fig. 4b). We also found that E-Cad::GFP levels were sometimes decreased locally at the base of these protrusions on the junctions (Fig. 4a, broken line). These observations suggest that actin protrusions can reduce the local levels of junctional E-cadherin. The junctions in Pak3-depleted cells also generated E-Cad::GFP-positive protrusions, but they were larger in size (Fig. 4a, arrowheads; Supplementary Movie 6). In addition, the local regions associated with E-Cad::GFP dilution on junctions were extended (Fig. 4a, broken lines). These observations suggest that Pak3 depletion enhances the reduction in E-cadherin levels at cell–cell junctions.

**Figure 4.**
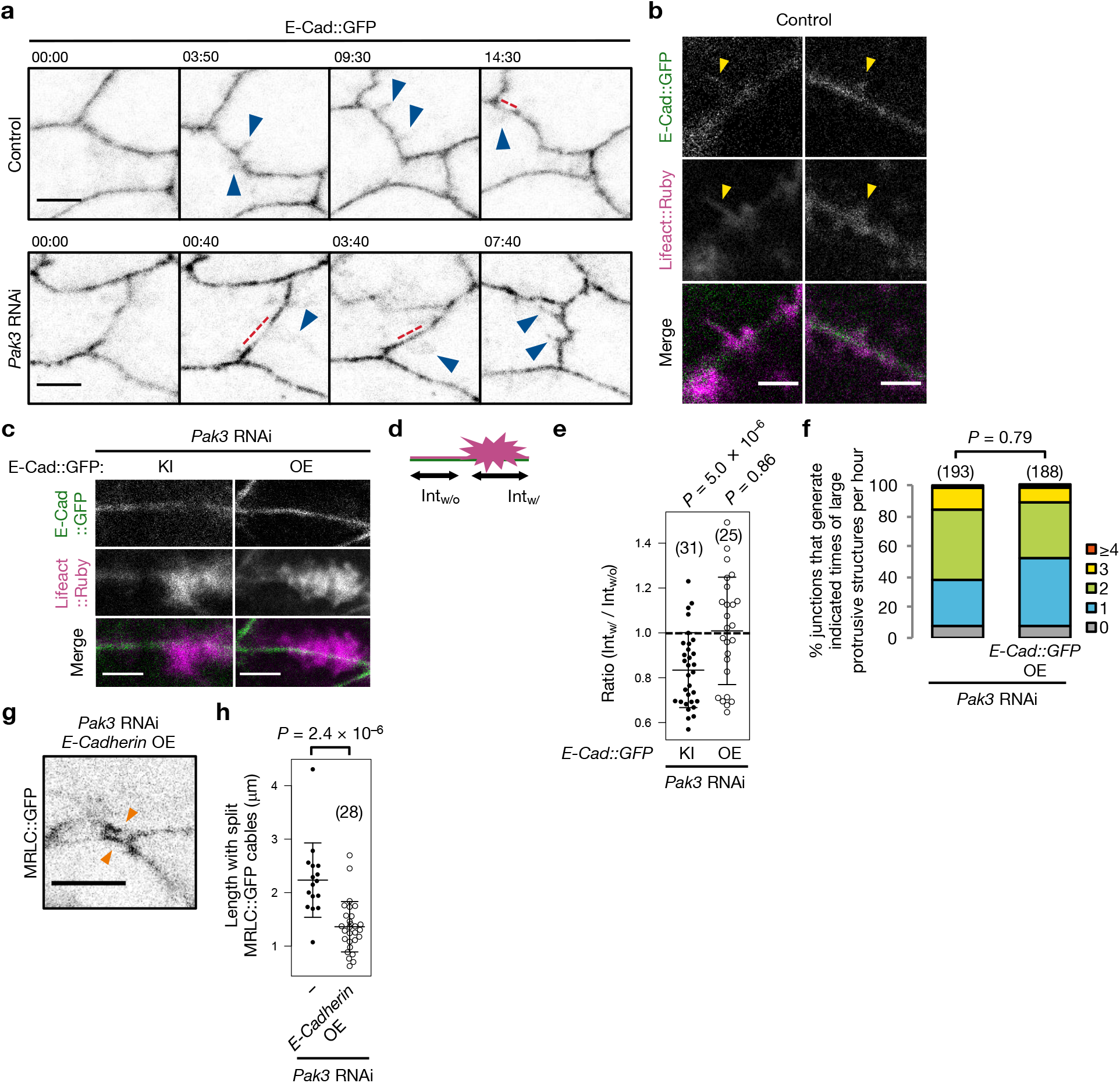
Reduction of E-cadherin levels at junctions induces myosin II dissociation. (**a**) Magnified time-lapse images of E-Cad::GFP at junctions. Arrowheads indicate E-Cad::GFP-positive protrusions, and broken lines indicate the local decrease in E-Cad::GFP levels at the bases of the protrusions. Scale bar, 3 μm. (**b** and **c**) Images of junctions with E-Cad::GFP and Lifeact::Ruby in control (b) and Pak3 RNAi (c) cells. Arrowheads indicate E-Cad::GFP and Lifeact::Ruby double-positive protrusions. Scale bar, 2 (b) and 3 (c) μm. (**d**) Schema of E-Cad::GFP intensities at local areas on the junctions with (Int_w/_) and without (Int_w/o_) large actin (magenta) protrusions. (**e**) Mean ± S.D. of the ratio of Int_w/_ and Int_w/o_ per junction. Parentheses indicate the number of examined junctions from 4 (Pak3 RNAi) and 3 (Pak3 RNAi and E-Cad::GFP overexpression) pupae. *P*-value by unpaired *t*-test. (**f**) Percentages of junctions forming large actin protrusions for the indicated number of times per hour are shown. Parentheses indicate the number of examined junctions from 4 pupae per genotype. *P-*values by Mann-Whitney *U*-test. (**g**) Representative images of split MRLC::GFP cables (orange arrowheads) at junctions. Scale bar, 5 μm. (**h**) Mean ± S.D. of the lengths of junctions with split myosin II cables. The Pak3 RNAi data (left) are the same as in Fig. 3g. Parenthesis indicates the number of examined junctions from 6 pupae. *P*-value by Dunnett’s test. Genotypes: (**a**) *+/Y;E-Cad::GFP;AbdB-Gal4, UAS-H2B::ECFP/+* (Control) and *+/Y;E-Cad::GFP;AbdB-Gal4, UAS-H2B::ECFP/UAS-Pak3 RNAi*; (**b**) *+/Y;E-Cad::GFP/UAS-LifeAct::Ruby;AbdB-Gal4/+*; (**c** and **e**) *+/Y;E-Cad::GFP/UAS-Lifeact::Ruby;AbdB-Gal4/UAS-Pak3 RNAi* and *+/Y;Ubi-E-Cad::GFP/UAS-Lifeact::Ruby;AbdB-Gal4/UAS-Pak3 RNAi*; (**f**) *+/Y; UAS-Lifeact::Ruby/+;AbdB-Gal4/UAS-Pak3 RNAi* and *+/Y; UAS-Lifeact::Ruby/Ubi-E-Cad::GFP;AbdB-Gal4/UAS-Pak3 RNAi*; (**g** and **h**) *MRLC^AX3^/Y;MRLC-MRLC::GFP/UAS-E-cadherin;AbdB-Gal4, UAS-H2B::ECFP/UAS-Pak3 RNAi*.

We then asked whether this dilution of E-cadherin is involved in the formation of aberrant actin protrusions and/or myosin II segregation. To this end, we upregulated E-cadherin levels in the Pak3-depleted A8a cells and examined if this manipulation suppressed these dynamics. A comparison of E-Cad::GFP intensities at the local regions with and without actin protrusions (Int_w/_ and Int_w/o_, respectively) within each junction in Pak3-depleted cells revealed that the levels of endogenous E-cadherin tagged with GFP (knock-in, KI) were significantly reduced to approximately 80% in the presence of actin protrusions (Fig. 4c–e). Overexpressing E-Cad::GFP under the control of the ubiquitin promoter, which slightly increased E-cadherin protein levels^42^, attenuated the local reduction in E-Cad::GFP levels upon the formation of actin protrusions (Fig. 4c,e). However, this manipulation did not attenuate the generation of large actin protrusions, indicating that E-cadherin dilution is located downstream from the formation of actin protrusions (Fig. 4f).

We next induced the overexpression of intact E-cadherin^43^. This manipulation suppressed the dissociation of myosin II from cell–cell junctions in Pak3 RNAi cells, which was indicated by a reduction in the length of junctions with split MRLC::GFP cables, similar to the effect of Abi RNAi (Figs. 3f,g and 4g,h). Taken together, these results suggest that the dilution of E-cadherin at cell–cell junctions induced by actin protrusions results in the inability of Pak3-depleted cells to retain myosin II at the junctions.

### E-cadherin maintains tissue dynamics

We finally explored whether overexpressing E-cadherin also restored tissue dynamics in Pak3 RNAi flies. E-Cad::GFP overexpression partially restored the junction-remodeling frequency in Pak3-depleted cells and slightly decreased the frequency of the shortening defect (Fig. 5a). In addition, this manipulation decreased the number of adult males with abnormal orientations of genitalia (Figs. 1b and 5b). These results suggest that the E-cadherin-mediated association of myosin II at junctions is required for completing junction shortening, leading to proper cell intercalation and epithelial cell movement.

**Figure 5.**
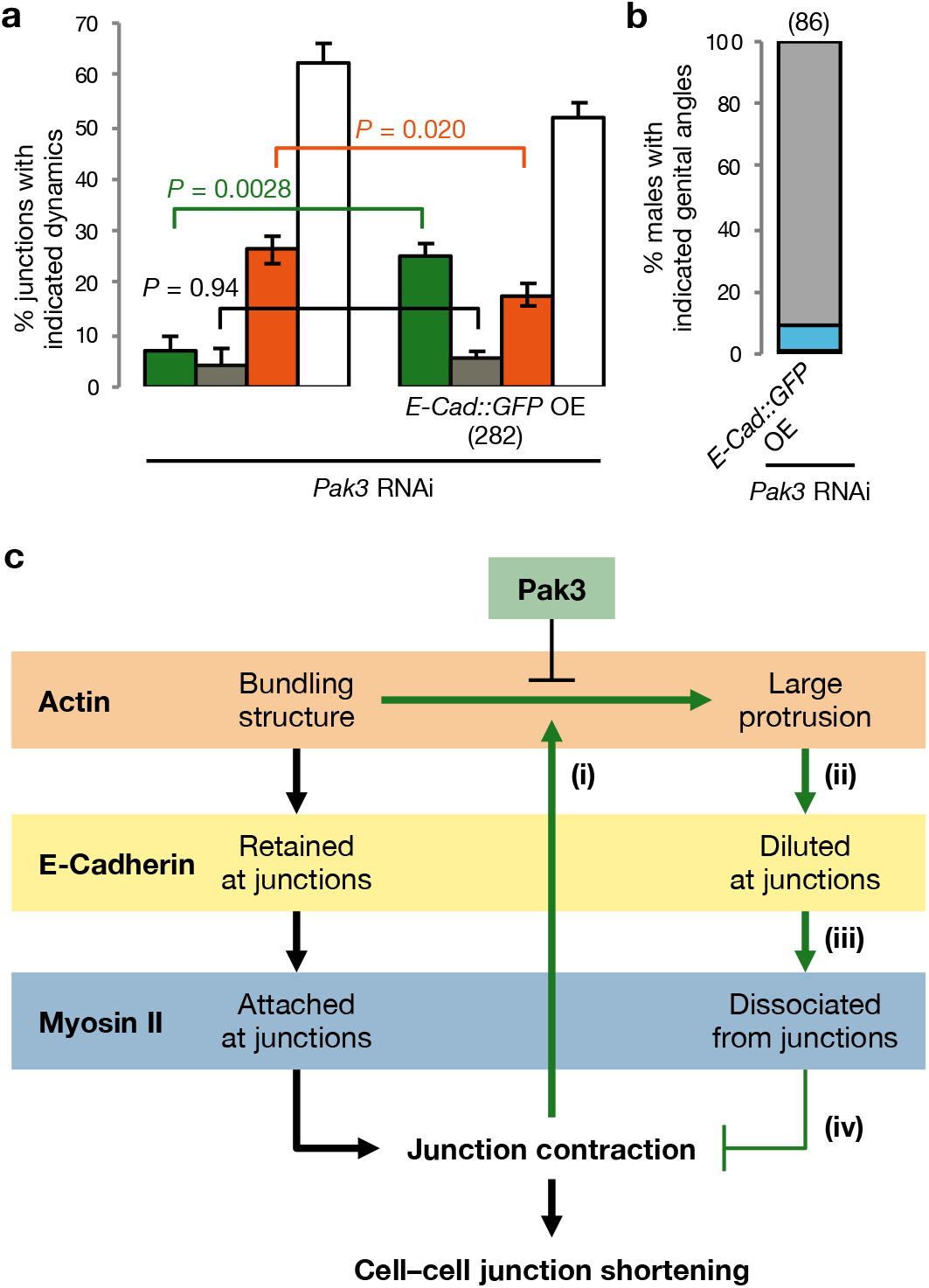
E-cadherin overexpression rescues tissue dynamics. (**a**) Mean ± S.D. of the percentages of junctions with the categorized dynamics. Bar colors correspond to Fig. 1e. The Pak3 RNAi data (left) are the same as in Fig. 1f. Parenthesis indicates the number of examined junctions from 3 pupae per genotype. *P-*values by Tukey’s test. (**b**) Percentages of male adult flies with genitalia angles indicated in Fig. 1b are shown. Parenthesis indicates the number of males examined. (**c**) Schema of Pak3-dependent junction contraction. (i)–(iv) indicate the negative feedback mechanism (green arrows, described in the Discussion section). Genotypes: (**a**) *+/Y;Ubi-E-Cad::GFP/E-Cad::GFP;AbdB-Gal4, UAS-H2B::ECFP/UAS-Pak3 RNAi*; (**b**) *+/Y;Ubi-E-Cad::GFP/+;AbdB-Gal4, UAS-H2B::ECFP/UAS-Pak3 RNAi*.

## Discussion

During cell–cell junction shortening, actin and myosin II accumulate at junctions, and the resultant actomyosin bundles generate contractile forces^5, 44^. In contrast, previous reports demonstrated that myosin II contractility potentiates actin unbundling and depolymerization in vitro and in vivo^45, 46^, which could compromise contraction. It has also been recently shown that cell–cell contacts concomitantly induce both myosin II-driven contraction and protrusive branched actin-mediated extension^10^. Therefore, it would be reasonable to suppose that there is a mechanism to maintain contractility for persistent cell–cell junction shortening. In this study, we found that, in the absence of Pak3, aberrant actin dynamics at junctions disturb junction contraction during their shortening. On the basis of our results, we suppose the following scenario in cells lacking Pak3 (Fig. 5c): (i) upon contraction of cell–cell junctions, actin dynamics at the junctions are altered to generate abnormally large protrusions; (ii) these aberrant protrusions cause the dilution of E-cadherin at the junctions; (iii) the decrease in E-cadherin levels weakens the association of myosin II cables at the junctions; and (iv) the dissociation of myosin II cables causes insufficient transmission of their contractility to the junctions, leading to a decrease in junctional tension and incomplete junction shortening. In the normal condition (in the presence of Pak3), Pak3 blocks excess formation of actin protrusions, keeps E-cadherin and myosin II localized at junctions, and thus ensures persistent junction contraction, which is necessary for completing cell–cell junction remodeling and the resultant tissue movement (Fig. 5c).

This scenario proposes a possible negative feedback mechanism (Fig. 5c, green arrows); the contractile forces of myosin II at junctions drive junction shortening, but they can oppositely perturb shortening by concomitantly altering actin dynamics, such as inducing the formation of WRC-dependent branched actin networks. Since Pak proteins are reported to regulate actin dynamics^47, 48^, Pak3 is a candidate for counteracting such undesirable actin dynamics, although its precise molecular mechanism is still unclear. Considering that Pak proteins and the WRC are activated by Rho GTPases^47, 49, 50^, it is possible that the presence of Pak3 sequesters these GTPases from the branching network-forming machineries. In addition, since Pak proteins are involved in a diverse array of biological events and have various substrates^31, 51–54^, the identification of Pak3 substrates in this context will further deepen our understanding of how cells accomplish persistent junctional dynamics.

## Methods

### Fly genetics

The following *Drosophila* stocks were used: *w^1118^*, *hs-flp*, *Act>CD2>Gal4*, *UAS-mCD8::mCherry*, *UAS-Lifeact::GFP*, *UAS-Lifeact::Ruby*, *His2Av::mRFP*, *UAS-Dicer2*, *UAS-MRLC* (*sqh*) *RNAi* (JF01103), and *UAS-Abi RNAi* (HMC03190) (Bloomington Drosophila Stock Center); *UAS-MRLC RNAi* (7916) and *UAS-Rock RNAi* (3793) (Vienna Drosophila Resource Center); *UAS-Pak3 RNAi* (14895R-1) (National Institute of Genetics, Japan); *UAS-E-cadherin* and *Ubi-E-cadherin::GFP* (Drosophila Genomics and Genetic Resources); *UAS-Pak3::GFP* (a gift from S. Hayashi); *AbdB-Gal4^LDN^* (ref. 24); *Pak3^d02472^* (ref. 30); *E-Cad::GFP*^28^; *sqh^AX3^;sqh-sqh::GFP* (MRLC::GFP)^55^; *sqh-UtrABD::GFP*^36^; and *UAS-Histone2B (H2B)::ECFP*^56^. The flies were raised, and all experiments were performed at 25°C. Somatic RNAi clones were induced using the FLP/FRT technique^57^ in white pupae (at 0 h APF) by heat shock (37°C for 15 min).

### Antibodies

Antibodies against Dlg (4F3; Developmental Studies Hybridoma Bank) and Pak3^58^ were used.

### Immunohistochemistry

Pupae were dissected and fixed in 4% paraformaldehyde in phosphate-buffered saline (PBS) for 20 min at room temperature (RT) and permeabilized with 0.1% Triton X-100 in PBS (PBT). The permeabilized samples were incubated in PBT with 5% donkey serum (blocking buffer) for 30 min at RT, incubated with primary antibodies in blocking buffer overnight at 4°C, washed with PBT, incubated in blocking buffer for 30 min at RT, and incubated with secondary antibodies (Alexa Fluor 647 goat anti-mouse IgG; Life Technologies) in blocking buffer for 2 h at RT. The samples were washed with PBT and mounted with 70% glycerol in PBS. Fluorescence microscopy images were captured on a TCS SP8 with a 63× numerical aperture (NA) 1.3 glycerol objective (Leica). Images are maximum intensity projections of serial optical sections taken at a 0.5-μm z step size.

### Live imaging

Pupae were prepared as described previously^21^. Time-lapse imaging of flies was performed using an SP8 confocal microscope with 63× NA 1.3 glycerol and 20× NA 0.75 dry objectives (Leica), except for the images in Figs. 1D and 3A and Movies S1 and S2, which were obtained using an inverted microscope with a 60× NA 1.3 silicone oil objective (Olympus) equipped with a spinning-disc confocal unit (CSU-W1; Yokogawa) and a Zyla 4.2 PLUS sCMOS camera (Andor). All images are maximum intensity projections at the level of the AJs taken at a 1-μm z step size, except for images showing rotation of the genitalia (Fig. 1A), which are maximum intensity projections of the posterior end of flies taken at a 5-μm z step size. Time-lapse images were acquired at 10-s, 20-s, 2-min, or 10-min intervals.

### Laser ablation

Laser ablation was performed with a 365-nm MicroPoint laser (Andor). To cut junctions, a 365-nm laser pulse was applied for 1 iteration to the point at the middle of the targeted junctions at the level of AJs, which was determined by maximum E-Cad::GFP intensity.

### Tracking of junction dynamics

Junction dynamics were analyzed manually with Fiji software. Projected time-lapse images were used. In the A8a epithelia, junctions were tracked from the initiation of movement (26 h APF) to the time when the genitalia angles were greater than 90° (30 h APF)^21^. Junctions that were remodeled at least once were categorized as “remodeling.” Junctions that failed to resolve four-way vertices were categorized as “growth defect.” Junctions that remodeled, but then immediately retracted the new junctions and re-formed in the original direction were also categorized as “growth defect.” Junctions that shortened to less than 1.5 μm in length, but failed to form four-way vertices were categorized as “shortening defect.” “No remodeling” included junctions that did not shorten to less than 1.5 μm in length.

### Quantification of junction length, fluorescence intensity, and actin dynamics

Cell–cell junction length was determined as the distance between vertices, except for when split MRLC::GFP cables emerged, which was determined as the distance between myosin II cables in cells surrounding the shortening junctions when the myosin II cables in cells forming the shortening junctions first segregated. The mean fluorescence intensity of E-Cad::GFP at a region on the junctions was measured using a line along the junctions. The frequency of the emergence of large actin protrusions was counted at each junction within 1 h.

### Statistical analysis

All statistical analyses were performed using R. To assess significance, the following tests were used: two-way analysis of variance (ANOVA) followed by Tukey’s test for comparing the percentage of remodeling junctions; two-way ANOVA followed by Dunnett’s test for comparing junction length with split myosin II cables among Pak3 RNAi cells; unpaired two-tailed *t*-test for comparing two samples; one-sample *t*-test for the significance of the Int_w/_/Int_w/o_ ratio with test values of 1; and unpaired two-tailed Mann-Whitney *U*-test followed by Bonferroni’s test for comparing the frequency of the appearance of actin diffusive structures.

## Acknowledgements

We thank S. Hayashi for providing the *UAS-Pak3::GFP* fly strain; N.T. Sherwood for providing the *Pak3^d02472^* fly strain; T. Lecuit for providing the *sqh-UtrABD::GFP* fly strain; N. Harden for providing the anti-Pak3 antibodies; E. Maekawa, A. Isomura and Y. Umegaki, S. Sato, K. Takahashi and A. Kuranaga for supporting the experiments; and members of the S. Hayashi, T. Nishimura, S. K. Yoo, Y.-C. Wang, and all member of our laboratory for discussions. We are grateful to “ThinkSCIENCE” for language proofreading of the manuscript. This work was supported by grants from JST CREST Grant Number JPMJCR1852, Japan (E.K.), the research grand for Astellas Foundation for Research on Metabolic Disorders (E.K.), the Takeda Science Foundation (E.K.), the Japan Foundation for Applied Enzymology (E.K.), MEXT KAKENHI grant number JP26114003 (E.K.), and JSPS KAKENHI grant numbers JP24687027 (E.K.), JP16H04800 (E.K.), and JP18K14691 (H.U).

## Author contributions

H.U. conceived the study, performed the experiments, analyzed the data, and wrote the manuscript. E.K. conceived the study, analyzed the data, and revised the manuscript.

## Competing interests

The authors declare no competing interests.

## Materials and correspondence

Further information and requests for resources and reagents should be directed to and will be fulfilled by the corresponding author, Erina Kuranaga.

---

**Supplementary Movie 1.** Dynamics of cell–cell junctions during the movement of control A8a cells. E-cadherin was visualized with E-Cad::GFP. Scale bar, 10 μm. Genotype, *+/Y;E-Cad::GFP;AbdB-Gal4, UAS-H2B::ECFP/+*.

**Supplementary Movie 2.** Dynamics of cell–cell junctions during the movement of Pak3 RNAi A8a cells. E-cadherin was visualized with E-Cad::GFP. Scale bar, 10 μm. Genotype, *+/Y;E-Cad::GFP;AbdB-Gal4, UAS-H2B::ECFP/UAS-Pak3 RNAi*.

**Supplementary Movie 3.** Dynamics of actin at cell–cell junctions in the control A8a cells. Actin was visualized with Lifeact::GFP. Scale bar, 10 μm. Genotype, *+/Y;UAS-Lifeact::GFP;AbdB-Gal4/+*.

**Supplementary Movie 4.** Dynamics of actin at cell–cell junctions in the Pak3 RNAi A8a cells. Actin was visualized with Lifeact::GFP. Scale bar, 10 μm. Genotype, *+/Y;UAS-Lifeact::GFP;AbdB-Gal4/UAS-Pak3 RNAi*.

**Supplementary Movie 5.** Dynamics of E-Cad::GFP at cell–cell junctions in the control A8a cells. Scale bar, 3 μm. Genotype, *+/Y;E-Cad::GFP;AbdB-Gal4, UAS-H2B::ECFP/+*.

**Supplementary Movie 6.** Dynamics of E-Cad::GFP at cell–cell junctions in Pak3 RNAi A8a cells. Scale bar, 3 μm. Genotype, *+/Y;E-Cad::GFP;AbdB-Gal4, UAS-H2B::ECFP/UAS-Pak3 RNAi*.

